# Refinement Strategies for Tangram for Reliable Single-Cell to Spatial Mapping

**DOI:** 10.1101/2025.01.27.634996

**Authors:** Merle Stahl, Lena J. Straßer, Chit Tong Lio, Judith Bernett, Richard Röttger, Markus List

## Abstract

**Motivation:** Single-cell RNA sequencing (scRNA-seq) provides comprehensive gene expression data at a single-cell level but lacks spatial context. In contrast, spatial transcriptomics captures both spatial and transcriptional information but is limited by resolution, sensitivity, or feasibility. No single technology combines both the high spatial resolution and deep transcriptomic profiling at the single-cell level without trade-offs. Spatial mapping tools that integrate scRNA-seq and spatial transcriptomics data are crucial to bridge this gap. However, we found that Tangram, one of the most prominent spatial mapping tools, provides inconsistent results over repeated runs.

**Results:** We refine Tangram to achieve more consistent cell mappings and investigate the challenges that arise from data characteristics. We find that the mapping quality depends on the gene expression sparsity. To address this, we (1) train the model on an informative gene subset, (2) apply cell filtering, (3) introduce several forms of regularization, and (4) incorporate neighborhood information. Evaluations on real and simulated mouse datasets demonstrate that this approach improves both gene expression prediction and cell mapping. Consistent cell mapping strengthens the reliability of the projection of cell annotations and features into space, gene imputation, and correction of low-quality measurements. Our pipeline, which includes gene set and hyperparameter selection, can serve as guidance for applying Tangram on other datasets, while our benchmarking framework with data simulation and inconsistency metrics is useful for evaluating other tools or Tangram modifications.

**Availability:** The refinements for Tangram and our benchmarking pipeline are available in https://github.com/daisybio/Tangram_Refinement_Strategies.

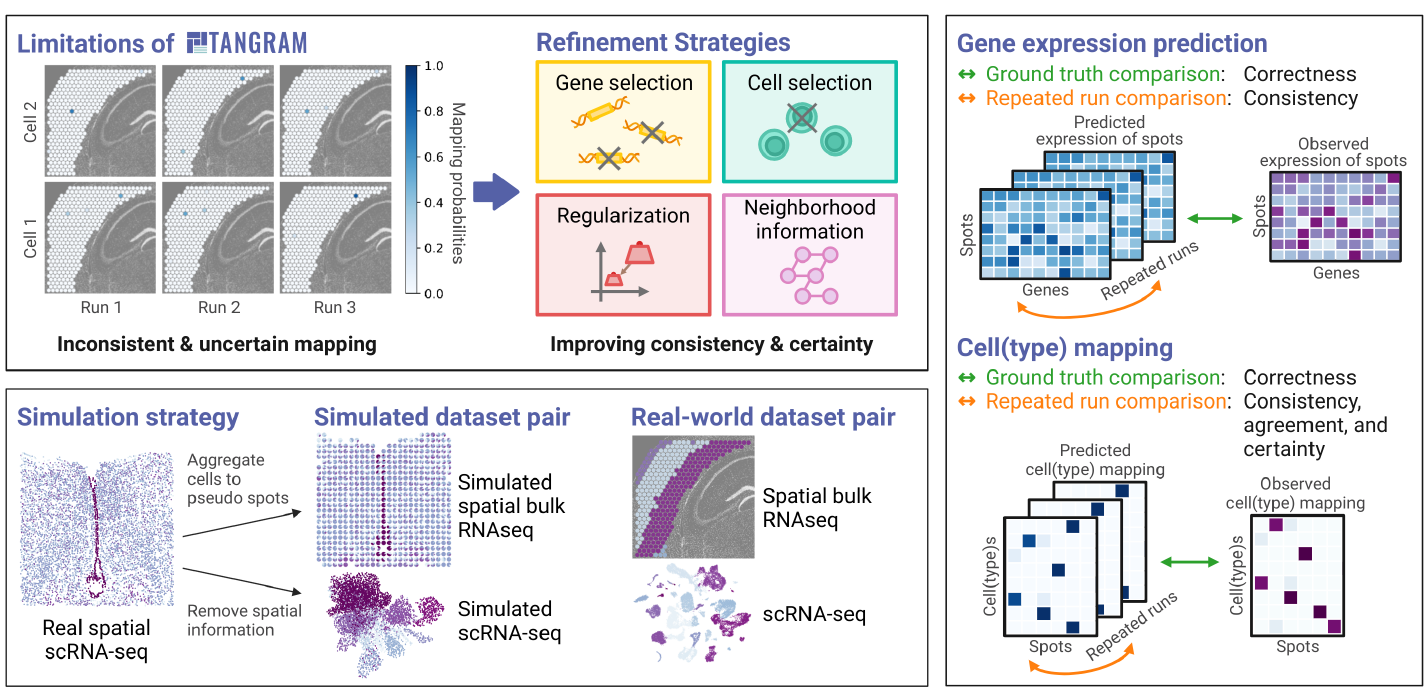

## 1. Introduction

Spatially resolving the transcriptome at the single-cell level is crucial for unraveling complex biological processes, including cell-cell communication, tissue organization, regeneration, niche formation, immune responses, tumor microenvironments, and differences between healthy and diseased tissues [Zhou et al., 2023, Sarkar et al., 2024, Singh, 2023, Vahid et al., 2023].

However, measuring such data remains a significant challenge due to limitations in current technologies. Single-cell RNA sequencing (scRNA-seq) provides detailed mRNA or, respectively, gene expression profiles at single-cell resolution but lacks spatial context. In contrast, recent advances in gene expression profiling methods allow for capturing the localization of cells and their associated gene expression within intact tissue samples, so-called spatial transcriptomics [Yue et al., 2023]. Spatial transcriptomics technologies vary widely in their strengths and limitations. In situ hybridization and sequencing technologies like multiplexed error-robust fluorescence in situ hybridization (MERFISH) [Chen et al., 2015] and sequential fluorescence in situ hybridization (seqFISH) [Lubeck et al., 2014] can capture spatial information at single-cell resolution but are often targeted and therefore restricted to detecting a limited number of genes [Yue et al., 2023]. In situ sequencing-based approaches like Visium [Ståhl et al., 2016] can achieve transcriptome-wide coverage, but often suffer from low capture efficiency and resolution [Piñeiro et al., 2022].

No spatial transcriptomics technology combines high spatial resolution and deep transcriptomic profiling at the single-cell level without tradeoffs in one or more areas like resolution, area size, sensitivity, transcriptome coverage, or cost. To overcome these challenges, computational tools have been developed to integrate scRNA-seq with spatial transcriptomics data, effectively combining the cellular resolution and deep transcriptomic profiling of scRNA-seq data with spatial context. In general, there are two main strategies to combine scRNA-seq and spatial data: deconvolution and mapping [Yue et al., 2023]. In deconvolution, we are interested in finding the cell type proportions of a spot consisting of more than one cell [Wei et al., 2022]. Instead, mapping focuses on the assignment of cells to spatial locations. This allows to project cell annotations like cell types and other modalities into space. Tangram [Biancalani et al., 2021] is a widely used and top-performing [Li et al., 2022, 2023] model for cell mapping, where its simple design makes it easy to incorporate and test the use of prior knowledge. As we show here, Tangram’s reliability is undermined by inconsistencies in its mapping performance across repeated runs, impacting the biological insights derived from the resulting data.

This study addresses and improves these inconsistencies. We propose a comprehensive set of metrics to evaluate Tangram’s correctness, consistency, certainty, and agreement across repeated runs. We evaluate and refine Tangram’s performance using both real scRNA-seq and Visium data, and simulated dataset pairs based on single-cell spatial technologies. We show that combining several refinement strategies leads to markedly improved cell mapping. In summary, we lay the groundwork for benchmarking cell mapping strategies and vastly improve the quality of spatially resolved single-cell data obtained with Tangram.

## 2. Material and Methods

We define the single-cell gene expression data matrix *S* with dimensions *n*_cells_ × *n*_genes_. Similarly, the spatial gene expression matrix *G* has dimensions *n*_spots_ × *n*_genes_. Using these, Tangram learns a *n*_cells_ × *n*_spots_ mapping probabilities matrix 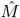. Furthermore, we denote the number of cell types in the single-cell dataset as *n*_cell types_ and the *n*_cells_ × *n*_cell types_ cell type one-hot-encoded matrix as *A*. This represents categorical cell type labels as a binary vector where all elements are zero except for a single one indicating the corresponding cell type.

### 2.1. Tangram’s mapping algorithm

Tangram [Biancalani et al., 2021] takes the single-cell matrix *S* and the spatial matrix *G* as input and predicts the mappings as a probability matrix 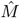 with 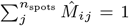 and 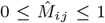 for all *i* ∈ [1, …, *n*_cells_] and *j* ∈ [1, …, *n*_spots_]. These constraints are enforced by applying the softmax function to an unadjusted, non-stochastic matrix 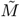. One of the main features of the mapping matrix is projecting genes, cell annotations, or other modalities *A* with the dimensions *n*_cells_ × *n*_annotations_ to spatial coordinates by computing 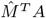. This enables predictions of gene expression or cell type mappings across spatial spots. The optimal mapping matrix minimizes the difference between the predicted spatial gene expression 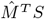, based on the mapping matrix and the single-cell matrix, and the observed spatial gene expression *G*. It can be computed using the following objective function based on the cosine similarity (CS)

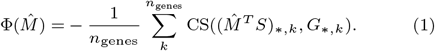

Gradient descent, in combination with backpropagation, is used for finding a minimum of this function. Equation 1 shows the default setting of Tangram, but the simplicity of the model also allows the incorporation of other terms to add, e.g., prior knowledge. Additionally, Tangram’s framework provides a constrained training mode. An *n*_cells_ long cell filter vector 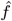 is introduced into the loss function such that only an optimal subset of the cells is mapped.

### 2.2. Tangram’s refinements

We evaluated Tangram’s performance across individual genes and cells. Based on observed insights, we developed modifications and extensions of Tangram, summarized in Figure 1.

**Fig. 1:**
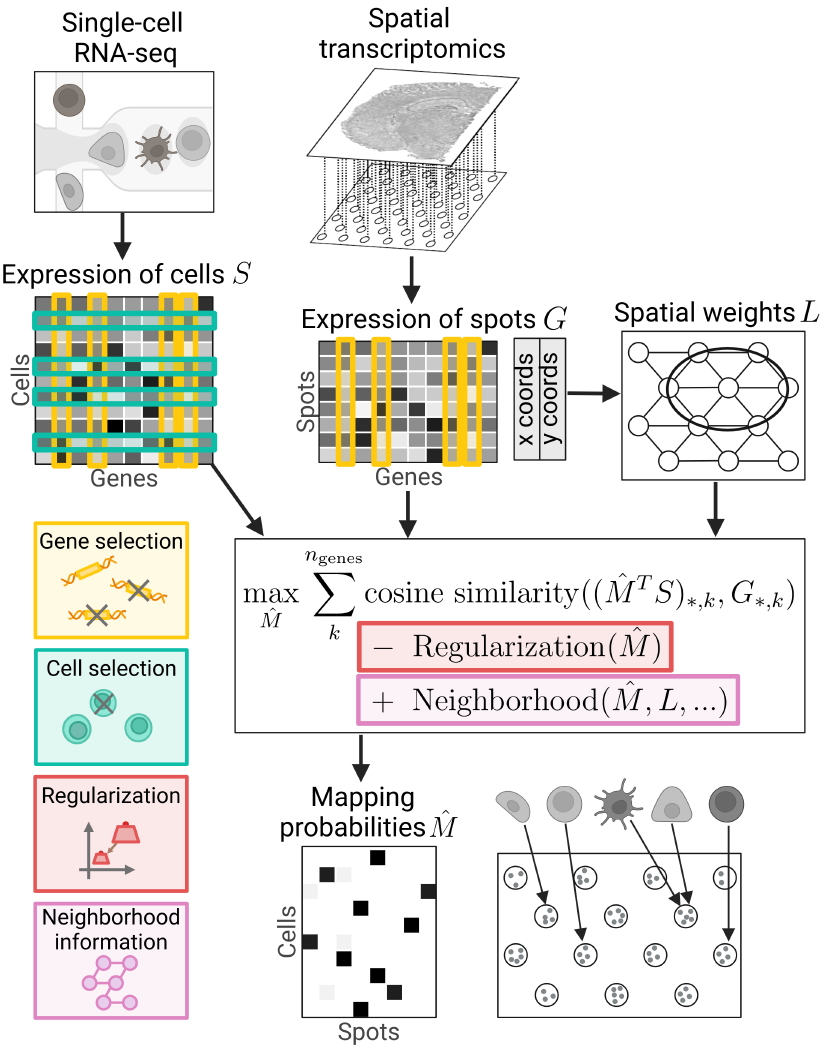
Overview of refinement strategies for Tangram. The options for gene selection (orange) and cell selection (green) are integrated. The objective function is augmented with regularization (red), and spatial context is incorporated through spot neighborhood information (purple).

#### 2.2.1. Gene selection

As Tangram training compares observed and predicted spatial gene expression, the gene set choice impacts results and runtime. As a baseline, we train the model on all available genes. Additionally, we test Tangram’s performance using an informative gene subset. To this end, we use the feature selection functionality of Spapros [Kuemmerle et al., 2024], a probe set selection method for targeted spatial transcriptomics based on scRNA-seq reference data. Since Spapros selects genes that capture both general and cell type-specific transcriptomic variation, we ensure a comprehensive representation of the original dataset.

#### 2.2.2. Regularization

Regularization is a widely used technique to prevent overfitting by introducing additional constraints to the optimization process.

Tangram offers entropy regularization to ensure that the probabilities of cell mappings are concentrated around a few spatial locations. In addition, we add an option for L2 regularization

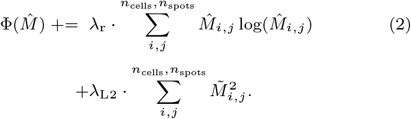

#### 2.2.3. Incorporating neighborhood information

Spatial information can be integrated in the form of *n*_spots_×*n*_spots_ spatial weight matrices *L* that capture the locality for each spot, including or excluding the spot itself. We add three extensions to the loss function that are based on *L*.

First, we apply spatial weighting to both the observed and predicted gene expression data of the spots before calculating the cosine similarity:

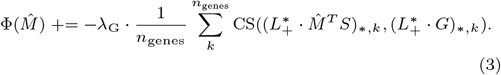

The *n*_spots_ × *n*_spots_ spatial weight matrix 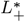 contains ones in the diagonal to include the spot itself and smaller fractions depending on the distances for the neighboring spots. This approach assumes that gene expression values for nearby spots are likely to be similar to each other. By applying spatial weights and essentially including gene expression data from neighboring spots, it smoothens the gene expression predictions, which allows for a certain degree of error for adjacent spots.

Second, we ensure that cells of the same type tend to cluster together in neighboring spots (cell type islands). The probability that a particular cell type maps to a given spot should be at least as large as the probabilities of that cell type mapping to the neighboring spots:

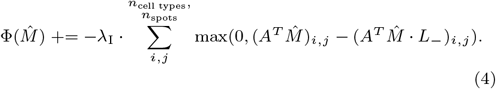

The *n*_cells_ × *n*_cell types_ one-hot-encoded matrix of cell types *A* is used to sum up the probabilities of all cells from one cell type for each spot. *L*_−_ is a spatial connectivity matrix (*n*_*spots*_ × *n*_*spots*_), which is used to collect the cell type probability sums from the neighboring spots. Its (*i, j*)-th entry is 1 if spots *i, j* are connected (*i* ≠ *j*), else 0. Of note, the spots are not connected to themselves to ensure that self-comparisons are excluded. This mechanism prevents spatial outliers and encourages spatial groupings of cells of the same type, reflecting biological reality.

A third method for incorporating spatial information is the use of local spatial indicators, which originate from geography. Local Indicators of Spatial Association (LISA) are designed to indicate the extent of spatial clustering of similar values around each observation [Anselin, 1995]. Thus, they perform a one-to-one mapping of the values. The local Getis-Ord *G*^∗^ statistic [Getis and Ord, 1992] measures spatial autocorrelation by identifying regions of high or low contrast. This spatial statistic is applied gene-wise to the spatial gene expression matrices. The goal is to maintain these statistical properties during the training process for the predicted gene expression based on the single-cell and the mapping matrix. This is achieved by extending the loss function with the term

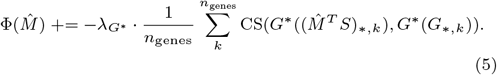

#### 2.2.4. Cell selection

Depending on the specific task and dataset, mapping only a subset of the cells may be beneficial. As mentioned, Tangram offers a constrained mode that allows the model to learn an optimal cell subset during training. However, this mode introduces additional variability across repeated runs, as the selected subset may differ each time. To address this when benchmarking, we use the constrained mode as a preprocessing step to identify a single, fixed cell subset that is used for every Tangram run. We estimate the number of target cells from a stained image where individual cells are visible, using a semantic segmentation method per the original Tangram study. Additionally, we explore an alternative cell sampling approach that is inspired by the CytoSPACE [Vahid et al., 2023] preprocessing steps. CytoSPACE uses estimations of cell type proportion and cell count number for each spatial spot to sample single-cell data that matches these counts. For comparability, we use a predicted cell filter generated with vanilla Tangram to supply these estimations.

### 2.3. Benchmarking datasets

To evaluate the performance of the original Tangram and the various modifications we propose, pairs of datasets consisting of single-cell gene expression data and spatial gene expression data are required. As suggested by Tangram’s developers, both datasets are normalized by the total counts over all genes per cell or spot and logarithmized before running Tangram. We test the models on one paired real-world scRNA-seq and Visium mouse cortex dataset pair. The scRNA-seq data contains 21,697 cells and 14,785 overlapping genes with the Visium counterpart, which consists of 324 spots.

Since the real datasets are missing the ground truth for the cell mapping, we generated low-resolution datasets based on data from spatial technologies with single-cell resolution, similar to [Sang-Aram et al., 2024], [Wei et al., 2022] and [Hao et al., 2024]. The gene expression data is summed up in a spatial grid, disregarding the spot-to-spot distance of the Visium technology. The bin size is adjusted according to the specific dataset to achieve average cell counts of 1-10+ cells like Visium spots. Sparse spots with less than 3 cells are excluded. The single-cell matrix is derived from the spatial single-cell dataset by removing the spatial information. We run the simulation based on a MERFISH mouse hypothalamus dataset and a SeqFISH mouse embryo dataset. The resulting simulated datasets comprise: (1) 6,185 cells, 589 spots in a 25 × 25 grid, and 161 genes, and (2) 19,416 cells, 1,688 spots in a 45 × 62 grid, and 351 genes.

Gene selection, as described in Section 2.2.1, is only applied to the Visium dataset since the MERFISH and SeqFISH datasets are already limited in gene number. Cell filtering (Section 2.2.4) is also not necessary for the MERFISH and SeqFISH datasets because our simulation produces one-on-one matched cells.

### 2.4. Benchmarking measures

Tangram is non-deterministic due to its random initialization. To quantify the consistency, we execute Tangram 10 times, each with a different random initialization. We evaluate the predicted spatial gene expression 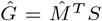, by comparing it across the repeated runs and against the ground truth *G*. Similarly, the predicted mapping matrix 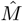 is assessed over repeated runs and, where available, compared to its ground truth *M*, which is a one-hot encoding of the cells belonging to a specific spot. Additionally, we calculate the *n*_cell types_ × *n*_spots_ cell type mapping prediction 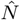 using the cell type one-hot-encoded matrix *A*. Following Tangram’s methodology, we normalize the summed probabilities to ensure they represent valid probabilities by dividing each entry in the matrix by the highest probability value for the corresponding cell type. This results in

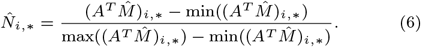

We call its ground truth *N* with

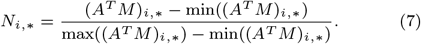

Again, we compare 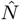 across repeated runs and, in the case of the simulated datasets, to the ground truth *N*.

#### 2.4.1. Measures for ground truth comparisons

##### Definition 1

The **gene expression prediction correctness** measures the similarity of the gene expression vectors for every gene 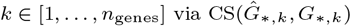.

##### Definition 2

The **cell mapping correctness** is defined as the difference of the predicted probabilities 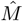 to the ground truth *M* and measured by the cross-entropy (CE) with 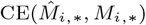 for every cell *i* ∈ [1, …, *n*_cells_].

##### Definition 3

To measure the **cell type mapping correctness** between the predicted 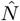 and ground truth mapping *N*, we extend this metric to a multi-label problem. Each probability is treated as a binary problem via the binary CE (BCE), where a cell type is either mapped or not mapped to a specific spot, and averaged over all spots with

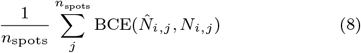

for every cell type *i* ∈ [1, …, *n*_cell type_].

#### 2.4.2. Metrics for comparing repeated runs

To see how similar the models perform over repeated runs, we run Tangram for *n*_runs_ = 10 times and compare their gene expression prediction and mapping probabilities. Suppose, 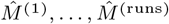 describe the different mapping probabilities predictions, and 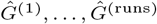 and 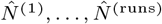 the corresponding gene expression prediction and cell type mapping probabilities.

##### Definition 4

The **gene expression prediction consistency** is measured via Pearson correlation (PC) with 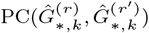 for all possible pairs 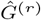 and 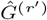 with *r, r*′ ∈ [1, …, *n*_runs_] and *r* ≠ *r*′ for every gene *k*∈ [1, …, *n*_genes_]. We consider the average PC of all possible pairs for each gene.

##### Definition 5

Similarly, we measure the **cell mapping consistency** for all cells *i* ∈ [1, …, *n*_cells_] and all possible pairs 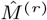 and 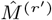 with *r, r*′ ∈ [1, …, *n*_runs_] and *r* ≠ *r*., via 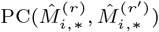.

##### Definition 6

In the same way, the **cell type consistency** 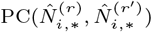 is computed for each cell type *i* ∈ [1, …, *n*_cell types_] and each possible pair 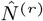 and 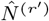 with *r, r*′ ∈ [1, …, *n*_runs_] and *r* ≠ *r*′. Again, we look at the mean PC over all possible pairs for each cell or cell type.

Since equally distributed probabilities tend to have a high Pearson correlation, we also evaluate the certainty and agreement of the mapping probabilities in similar ways to what is used for disagreement sampling [Danka, 2018].

##### Definition 7

To compute a vote cell mapping over repeated runs, we select for each run and each cell the spot with the highest probability as a vote and take the distribution over those with

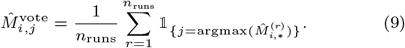

The 𝟙 is the indicator function with value 1 if the condition holds and 0 otherwise.

Applying the entropy (E) on the vote mapping (vote entropy) with 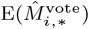 measures the certainty of the vote mapping and, therefore, also evaluates the **cell mapping agreement** over repeated runs.

##### Definition 8

Since the cell type mapping is a multi-label problem, the vote cell type mapping is computed by selecting all probabilities larger than 0.5

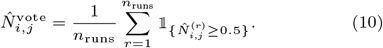

The **cell type mapping agreement** is evaluated by the cell type vote entropy via binary entropy (BE):

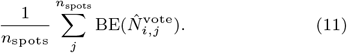

In this way, we can evaluate to what degree repeated runs agree with each other in selecting spots for the cells or cell types.

##### Definition 9

The consensus cell mapping for all repeated runs is the average of the class probabilities with

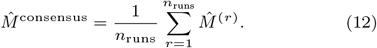

The consensus entropy 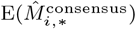 measures the certainty of these average probabilities and hence especially the **cell mapping certainty** over all runs.

##### Definition 10

To measure the **cell type mapping certainty**, this is extended to a multi-label problem by treating each probability as a binary problem and averaging over all spots with

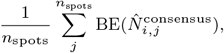

where 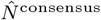 is defined in accordance to Equation 12.

If not already provided, we normalize the measures for correctness, consistency, agreement, and certainty to a range between 0 to 1 to enhance comprehensibility. This is achieved by dividing the entropy values by either log(*n*_spots_) or log(2) in multi-label scenarios, and by scaling the cross-entropy using the largest observed value. Furthermore, we take the counter value for the (cross-)entropies with 1 − *x* for *x* ∈ [cross-entropy, vote entropy, consensus entropy], such that 1 indicates a good and 0 a bad result for all measurements.

## 3. Results and Discussion

### 3.1. Impact of data properties on Tangram’s performance

To identify Tangram’s limitations, we analyzed its performance against dataset characteristics from the perspectives of genes and cells. Tangram was trained using default parameters and all overlapping genes on the real-world mouse cortex dataset pair. Its performance in predicting spatial gene expression was assessed against the ground truth and evaluated for consistency across 10 repeated runs. Additionally, we examined the consistency, agreement, and certainty of cell and cell type mappings across the repeated runs.

#### 3.1.1. Sparsity impacts gene expression prediction and overfitting

In the first step of the analysis, we focused on evaluating the performance of Tangram by measuring the correctness and consistency of gene expression prediction for individual genes. We trained Tangram on a 85% random training gene set and left out a 15% test gene set. We investigated the two metrics for both gene sets in relation to key data characteristics such as the number of cells and spots from both datasets where the gene is expressed.

This analysis reveals a dependency between the correctness of predictions and the sparsity of the data (Figure 2A). Genes expressed in fewer cells or spots tend to have poorer predictions. This observation aligns with observations of the Tangram study [Biancalani et al., 2021], which demonstrated that almost all non-sparse genes are predicted correctly with a large cosine similarity, while sparse genes often suffer from prediction errors caused by technical dropouts. Spatial data is typically more sparse than scRNA-seq data. In such cases, Tangram struggles to align gene expression patterns accurately due to uneven dropout rates between datasets. However, it has been argued that the prediction based on the more comprehensive single-cell data is more accurate than the measured expression in a spatial assay, and, therefore, Tangram can impute and correct spatial low-quality measurements.

**Fig. 2:**
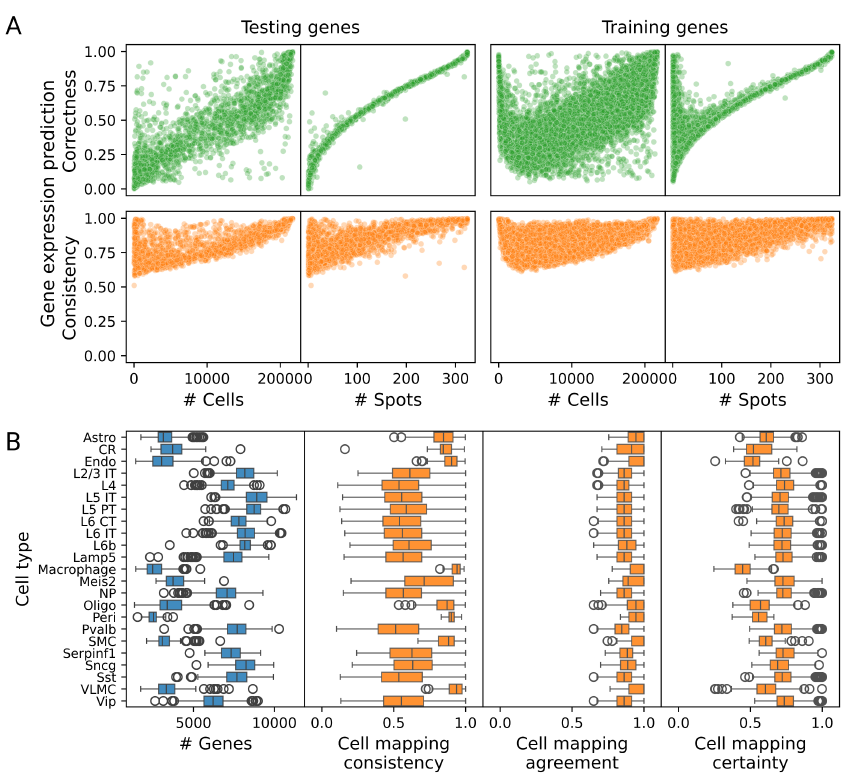
Data properties impact Tangram’s performance. (A) Gene expression prediction performance metrics on test and train gene sets are plotted against data sparsity measures. (B) Cell mapping performance metrics are visualized stratified by cell types.

The consistency of the predicted gene expression shows a similar pattern, where sparse genes were predicted less consistently than non-sparse genes.

Interestingly, Tangram shows a surprisingly good prediction performance for sparse training genes compared to the sparse genes in the test set. This discrepancy suggests overfitting, which is particularly problematic due to the difference in dropout rates between the datasets.

#### 3.1.2. Consistent but uncertain mapping of gene-sparse cells

Including dataset properties and cell type annotations from the single-cell dataset in the benchmarking reveals how data sparsity influences Tangram’s performance for individual cells.

We assessed characteristics of the single-cell data, alongside benchmark metrics for individual cells stratified by their cell types (Figure 2B). We refer to cells with a low number of expressed genes as gene-sparse cells. Cell types like astrocytes and Cajal-Retzius cells (CRs) stand out as they contain a large number of gene-sparse cells. Evaluating their mappings shows a large consistency and agreement over repeated runs. However, their certainty score is lower, meaning that the probabilities are more evenly distributed across the spatial locations, reducing the model’s confidence. On the other side, less gene-sparse cells show room for improvement as well, particularly in their consistency metrics.

### 3.2. Learnings and Tangram’s refinements

Based on the results from the analysis of Tangram’s performance, we developed several improvement strategies.

The gene expression prediction results show that Tangram struggles with sparse genes, where performance is worse regarding correctness and consistency. Sparse genes often have higher dropout rates in spatial data than single-cell data, making it harder for Tangram to accurately predict gene expression patterns. Additionally, overfitting to these sparse training genes exacerbates these issues, creating a trade-off between achieving high gene expression prediction accuracy and ensuring correct cell mappings. This trade-off stems from Tangram’s assumption that gene expression patterns perfectly correlate between single-cell and spatial datasets, which is unlikely to hold in real-world datasets. As a result, focusing too much on gene expression accuracy can lead to models that overfit to dataset-specific errors rather than capturing true biological patterns. These findings highlight the need for simulated datasets with cell mapping ground truths, as well as the limitations of using cosine similarity alone for training, and point to the need to incorporate additional information or refinements for Tangram.

First, we tested using informative gene subsets focusing on genes with significant biological relevance and lower sparsity. Furthermore, we assessed several regularization techniques to address the problem of overfitting.

We observed that sparse cells show higher consistency and agreement but suffer from reduced certainty in their mappings. This means that the model predicts lower probabilities more evenly across the spatial locations, which increases the consistency values and lowers the model’s confidence in pinpointing exact locations. However, higher agreement values suggest that the same spot consistently holds the highest probability across repeated runs, indicating that the model may have some understanding of the cell’s position. Sparsity can arise either from biological factors or technical dropouts, where the latter can provide the model with misleading information. Regardless of the cause, the lack of informative features makes it challenging for the model to learn effectively. Additionally, some cells are inherently less specific to particular regions and can naturally occur across multiple areas. These cells present further challenges, as their distribution may reflect true biological variability rather than technical limitations or model inaccuracies. Less sparse cells show room for improvement as well, especially regarding consistency.

To address this, we incorporate neighborhood information during training. Moreover, it might be beneficial to consider a cell subset mapping, where excluding, for example, sparse cells could help achieve a better overall mapping result. We benchmarked Tangram with its built-in cell filtering during training and the extensions, including a cell filter preprocessing step, to ignore the variability of the cell selection step. While filtering out sparse or poorly performing cells can work well for large cell types without significantly affecting their mapping, it poses a significant challenge for rare cell types. When only a few cells are measured for a rare type, and those cells are sparse, excluding them risks entirely compromising the mapping for that cell type. Therefore, we included a cell sampling preprocessing step based on the estimated cell number per cell type from the spatial dataset in our analysis.

### 3.3. Benchmarking Tangram refinement strategies

We trained each model 10 times on the real and the two simulated datasets and evaluated its performance regarding gene expression prediction, cell mapping, and resulting cell type mapping.

We tested the default Tangram setting trained on all possible genes as reference and performed a one-sided Mann-Whitney *U* test against it for every model. Additionally, a random assignment of cells to spots is used as baseline to demonstrate how the different metrics perform.

#### 3.3.1. More reliable cell mappings based on informative gene subsets

We evaluated Tangram trained on (1) highly variable genes from the single-cell data, (2) cell type-specific genes, (3) spatially variable genes, (4) the union of (1)-(3), and (5) Spapros genes. We found that Spapros performed best across all measures and, therefore, omitted the others.

As shown in Figure 3, Tangram trained on Spapros genes significantly outperformed the baseline vanilla model in terms of consistency, agreement, and certainty metrics, while maintaining comparable or slightly lower levels of gene expression prediction correctness. Additionally, this approach reduced computation time for training by a factor of 4.2, compared to training on all genes (35.8 s vs 8.6 s on two AMD EPYC 7313 16-Core Processors and one NVIDIA A40 64-Core GPU).

**Fig. 3:**
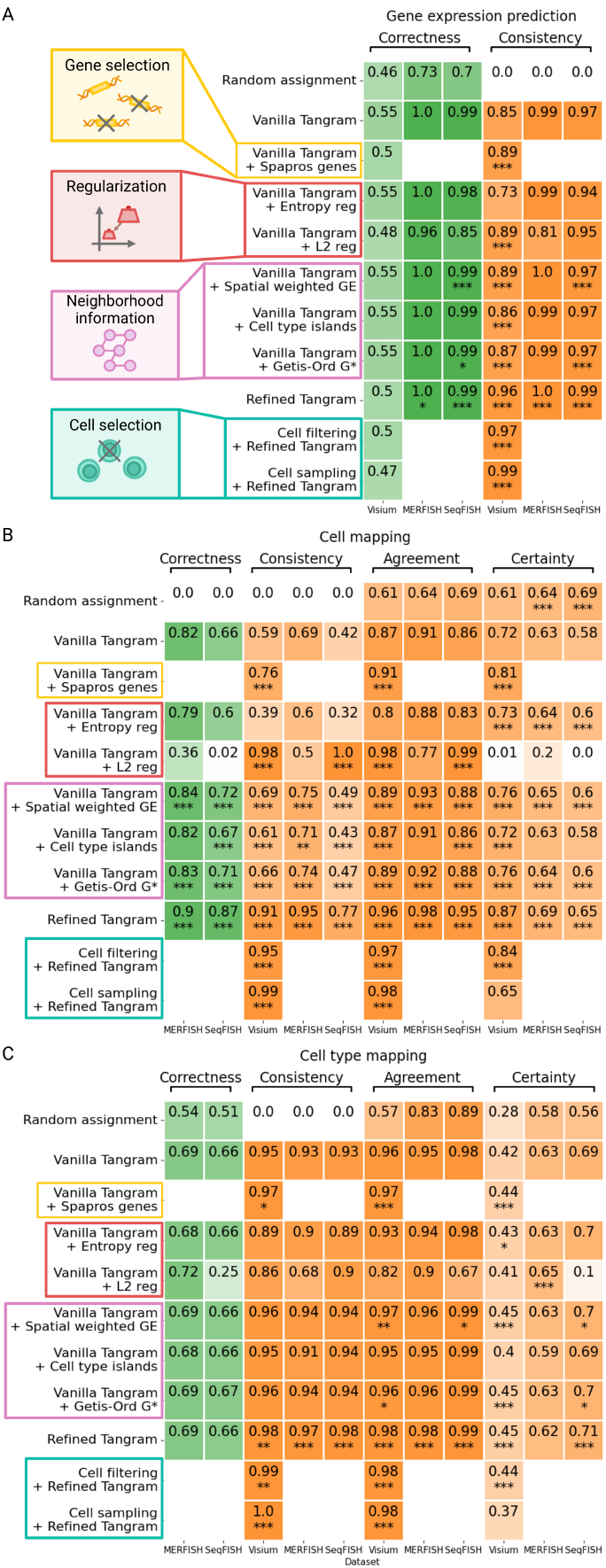
Benchmarking vanilla Tangram and refinement strategies. We mapped cells of real-world and simulated dataset pairs using vanilla Tangram, different refinement strategies, and a refined version of Tangram that integrates all beneficial refinement strategies. A random assignment of cells to spots is included as baseline. Metrics for ground truth comparisons are highlighted in green and metrics for repeated run comparisons in orange. A one-sided Mann-Whitney *U* test against the vanilla model was performed, where a p-value < 0.05 is indicated by ^*^, < 0.01 by ^**^, and < 0.001 by ^***^.

#### 3.3.2. Regularization: balance between certainty and consistency

We tested the integration of multiple regularization strategies in the objective function of Tangram.

As shown in Figure 3, entropy regularization improved certainty values by encouraging more confident predictions but reduced consistency. Including an L2 regularization term in the loss function improved consistency and agreement in cell mappings but significantly lowered certainty, making predictions less confident.

These results reveal a key trade-off between consistency and certainty when using regularization in Tangram. Finding the right balance between these two is crucial for achieving the best mapping results.

#### 3.3.3. Neighborhood information contributes to more reliable cell mappings

We evaluated various approaches, including spatial weighting of gene expression (Equation 3), enforcing cell type islands (Equation 4), and preserving local spatial indicators using local Moran’s *I* statistic [Moran, 1950], local Geary’s *C* statistic [Geary, 1954], and the local Getis-Ord *G*^∗^ statistic (Equation 5). Among these statistics, the local Getis-Ord *G*^∗^ statistic, with its differentiation between cold and hot spots, proved most effective. Followed by spatial weighting and cell type islands. Therefore, we include these three strategies in our analyses.

Adding spatial information through these approaches significantly improves performance across all metrics for cell mapping and gene expression prediction (Figure 3A and B). For cell type mappings, the methods lead to slight improvements, especially in agreement and certainty (Figure 3C). The biggest gain was seen in cell mapping consistency, with increases of up to 0.1, without negatively impacting other metrics.

#### 3.3.4. Combining modifications achieves the best cell mappings

We integrated all proposed modifications and guidelines to develop an optimal model, utilizing hyperparameter tuning on the real dataset pair using the Spapros gene set. The objective is to maximize the cell mapping consistency, agreement, and certainty while also ensuring high correctness of gene expression predictions, assessed via a 15% validation gene set.

The evaluation of the refined Tangram model demonstrates significant improvements in overall performance (Figure 3).

For gene expression prediction, the combined model shows significant yet relatively small improvements in consistency and correctness for simulated datasets. It is important to note that these metrics were already near their maximum values in the vanilla model. In contrast, the real dataset shows a minor reduction in gene expression correctness, which may be advantageous because of the trade-off between gene expression prediction and actual mapping correctness that stems from discrepancies between single-cell and spatial assays.

Our refinements result in substantial and consistent improvements across all cell mapping metrics. The most significant improvement is observed in cell mapping consistency, which addresses a key limitation of the vanilla model. On the real dataset, consistency increases markedly from 0.59 to 0.91 without cell subset selection and further to 0.99 when subset selection using filtering or sampling is applied during preprocessing. However, it is worth noting that cell subset selection is associated with reduced certainty values. Similar improvements are observed in simulated datasets, with increases of 0.26 and 0.35.

Cell type mapping performance remains comparable to the vanilla model with only small but significant improvements, particularly for the real dataset.

Our refined Tangram demonstrates comparable performance in gene expression prediction and cell type mapping, with small but significant increases. The most notable improvements are achieved in cell mapping performance, with large gains across all metrics, particularly consistency. These results indicate that the refinements effectively address the limitations of the baseline model, especially in enhancing the consistent differentiation between individual cells. As a result, the refined model offers more reliable spatial cell mappings.

## 4. Conclusion

In conclusion, we show that straight-forward, gradient-descent-based mapping of single-cell to spatial gene expression data achieves reliable results only if neighborhood information and regularization are included in the optimization process and the data preprocessing includes gene and cell filtering.

Comparing cell-to-location mappings computed by Tangram, we observed striking differences between repeated runs, which limits the tool’s reliability. Using both real and simulated datasets, we systematically assessed Tangram’s performance and identified critical areas for improvement.

Addressing these shortcomings, we proposed several refinements, including (1) optimizing gene set selection through approaches like Spapros, (2) employing regularization techniques to balance consistency and certainty, (3) incorporating spatial information using, e.g., neighborhood-based indicators, and (4) testing strategies for improved cell subset selection. Combined with hyperparameter tuning, these modifications significantly improved performance across all evaluated metrics, enhancing Tangram’s reliability for single-cell spatial mapping. While our refinements improved Tangram’s performance, limitations remain. The benchmarking framework could benefit from alternative spatially weighted metrics and more diverse datasets reflecting realistic biological complexities. Additionally, relying on idealized datasets and simulations necessitates further validation across more challenging experimental datasets, including those with higher cell type diversity and noise. Future research should expand the scope of benchmarking to include other cell mapping tools, providing a comparative landscape of reliability. Moreover, advancing Tangram’s application to three-dimensional datasets, such as those generated by MERFISH, could open new avenues for spatially resolved studies. Integrating prior information, such as ligand-receptor interactions and exploring embeddings or ensemble approaches, could further enhance mapping robustness. The presented refinements not only improve Tangram’s application for mapping scRNA-seq data to spatial contexts but also provide a foundation for projecting other modalities, such as chromatin accessibility and multi-omics datasets, into space. As spatial transcriptomics technologies evolve, the flexibility of Tangram’s architecture and its adaptability to emerging datasets make it a valuable tool for the field. By addressing fundamental challenges in cell mapping, this work contributes to more reliable single-cell spatial analyses, supporting the generation of comprehensive datasets and advancing our understanding of complex tissues.

## 5. Data and Code Availability

The code of the adapted Tangram algorithm and the benchmarking pipeline are available in https://github.com/daisybio/Tangram_Refinement_Strategies. We only used publicly available datasets that are, at the time of this publication, accessible via Squidpy v.1.6.2 [Palla et al., 2022].

- MERFISH mouse hypothalamus: We selected bragma 9 from the MERFISH dataset of [Moffitt et al., 2018].
- SeqFISH mouse embryo: The dataset was published in [Lohoff et al., 2020].
- Visium and scRNA-seq mouse cortex: The scRNA-seq data from [Tasic et al., 2018] was reduced to cortex layers. The spatial counterpart was taken from 10x Genomics Visium H&E [10x Genomics, 2020].

## 7. Author contributions statement

M.L. and C.L. conceived and designed this study. J.B. identified the limitations of Tangram. M.S. implemented and executed the algorithm adaptations and benchmarks. M.S. and L.S. interpreted the results and drafted the manuscript. M.L. supervised the project. All authors provided critical feedback and contributed to the final manuscript.

## 8. Acknowledgments

Funded by the Deutsche Forschungsgemeinschaft (DFG, German Research Foundation) [422216132]. This work was supported by the Novo Nordisk Foundation as part of the Data Science Collaborative Research Programme 2022 [NNF22OC0076414 (MOPITAS)]. We acknowledge support from the Munich Data Science Institute (MDSI) at the Technical University of Munich (TUM) via the Linde/MDSI Doctoral Fellowship program. We thank Konstantin Schröder for advice on the benchmarking measures and the statistical test. We thank Julian Owezarek for his help in deriving Equation 4. All figures are created in BioRender. Straßer, L. (2025) https://BioRender.com/e68m527

